# Biological screens from linear codes: theory and tools

**DOI:** 10.1101/035352

**Authors:** Yaniv Erlich, Anna Gilbert, Hung Ngo, Atri Rudra, Nicolas Thierry-Mieg, Mary Wootters, Dina Zielinski, Or Zuk

## Abstract

Molecular biology increasingly relies on large screens where enormous numbers of specimens are systematically assayed in the search for a particular, rare outcome. These screens include the systematic testing of small molecules for potential drugs and testing the association between genetic variation and a phenotype of interest. While these screens are “hypothesis-free,” they can be wasteful; pooling the specimens and then testing the pools is more efficient. We articulate in precise mathematical ways the type of structures useful in combinatorial pooling designs so as to eliminate waste, to provide light weight, flexible, and modular designs. We show that Reed-Solomon codes, and more generally linear codes, satisfy all of these mathematical properties. We further demonstrate the power of this technique with Reed-Solomon-based biological experiments. We provide general purpose tools for experimentalists to construct and carry out practical pooling designs with rigorous guarantees for large screens.

## Significance

We provide a mathematical formulation of biologically relevant structures in combinatorial pooling designs. Before this work, biologists had not articulated in precise ways the type of structures necessary or useful in pooling designs. We also show that Reed-Solomon codes, and more generally linear codes, satisfy all of these mathematical properties. We establish fresh connections between biologically relevant and combinatorial properties of error correcting codes and how to construct practical, relevant pooling designs with rigorous guarantees.

## Introduction

The field of molecular biology increasingly relies on large screens where overwhelming numbers of specimens are systematically assayed in the search for a particular outcome. These screens can come in various forms: systematic testing of tens of thousands of small molecules in a search for potential drugs; interfering with the activity of every gene in the genome to elucidate the organization of genetic networks that underly cancerous processes using RNAi; determining the ability of one protein to bind to every other protein in the genome in yeast two-hybrid systems; and testing the association between virtually every possible genetic variation in the genome and some phenotype of interest such as diabetes or heart diseases. One advantage of these screens is that they are considered “hypothesis-free,” letting the data reveal new and unexpected outcomes that are hard to deduce from current biological knowledge. The downside is that screens can be quite resource and energy consuming. In many cases, screens “waste” an overwhelming number of assays that deliver negative and uninteresting results to find the scarce positive findings.

In the last few years, we and others have independently demonstrated screens in various biological domains that are based on combinatorial pooling (see the Table in the supplementary material). These experiments have two basic commonalities. First, instead of independently assaying each one of the specimens, the experiments begin with grouping the specimens into pools using specific mathematical rules, and then assaying the pools. This procedure dramatically reduces the costs of the experiments as the number of pools is much smaller than the number of the original specimens. Second, in contrast with naive pooling in which a group of specimens participate in exactly one pool, each specimen in combinatorial pooling participates in a series of unique pools. Thus, it is possible to assign a unique result to each specimen by comparing the pattern of the results to the pooling design, as long as the number of positive findings is relatively small.

Most of the experiments above exploited **non-adaptive group testing** or **compressed sensing** to design the pooling pattern. These two closely related mathematical domains deal with construction of efficient designs that reduce the number of pools while preserving the ability to assign unique results to each item. Under these frameworks, biological screens are cast as a system of (linear) equations

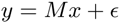

with *n* variables, each variable corresponds to an item in the screen. The items can come in different forms: small molecules in chemical screens, different hairpins in RNAi screens, a library of preys in yeast two-hybrid screens, or human individuals in genetic studies. The aim of the screen is to uncover *x*, a vector of length *n*, whose *i*th position denotes the biological property of item *i*. For example, in a small molecule screen, *x* can be either 1 or 0, where 1 denotes an active compound and 0 denotes an inactive compound. The experimenter does not observe *x* directly. Rather, she employs *t* assays into which the *n* items have been pooled and observes *y*, a vector of length *t* whose *j*th position denotes the assay results for *j*th assay. For example, in a small molecule screen, the entries of *y* can denote the fraction of surviving cells in the presence of the compound, or *y* can be a vector with entries that are either 0 or 1, where 0 means that no cell survives while 1 captures the survival of some of the cells in the assay. Finally, we can express the pooling design by *M*, a matrix with *t* rows and *n* columns whose entries are non-negative numbers. The entry *M_ij_* = *r* means that *r*-units (micrograms, microliters, moles, or some arbitrary measure) of item *j* are assayed in the *i*th reaction. With this representation, trivial screens (no pooling) simply correspond to the *n* × *n* identity matrix; each items is assayed in exactly one distinct reaction. The noise *ϵ* can be zero, a random variable, or an arbitrary, bounded value, depending on the model. In group testing, the product *Mx* is computed using Boolean arithmetic: the result of an assay is positive if and only if at least one sample in the pool is positive. In compressed sensing, the entries of the matrix *M* are real numbers and the product *Mx* is computed using standard real-valued arithmetic.

Unfortunately, the theoretical work in group testing and compressed sensing does not address many practical design challenges biologists face in using such pooling strategies. Significant progress has been made in developing pooling designs with a minimal number of assays but with scant attention to whether these designs can be implemented in practice.

Here, we map the current implementation gaps and propose a mathematical framework based on coding theory that addresses most of these barriers. We also describe experimental validations of Reed-Solomon code and provide an overview of web-based tools that we have developed for creating and implementing pooling designs at the bench. Using these tools, experimentalists with no mathematical knowledge in coding theory and minimal equipment can conduct complex pooling experiments.

## Challenges and Results

**Features of practical pooling designs**. In the last few years, we have conducted dozens of large scale pooled screens in various biological domains that highlight common features and issues in implementing pooled experiments.

***Well-balanced designs***. We noticed that the most implementation-robust designs are well-balanced designs, where each item is pooled exactly the same number of times and all the assays have (close to) an identical number of items. These designs consume the same amount of material from all items, allowing straight forward planning of the experiment and reducing the risk that an item will run out during pooling. It also ensures that each item is treated equally. Assays with identical numbers of items are also highly beneficial. They produce more consistent results that mitigate the effect of diluting the specimens. Another by-product is that well-balanced designs allow fast quality assessment of the pooling procedure before starting the expensive screening step. Large deviations in pool volumes or in the residual specimen material can be easily detected by eye and serve as an indicator that there was a potential flaw in constructing the pools.

***Maximal utilization of biological kits***. A large number of biological kits are sold in batches of 96 assays to fit the common microtiter plates. In many realistic scenarios there are minimal cost differences between a design with 50 assays or 96 assays, but a significant difference between a design with 96 pools and 97 pools. Thus, it is desirable to find a pooling design with a multiple of 96 assays. Of course, one can construct a random pooling design to achieve this aim, but then the design will not be well-balanced. Thierry-Mieg [22] proposed the Shifted Transversal Design (STD), which is a well-balanced and was widely used in biological experiments. Unfortunately, STD cannot fully utilize a 96 well plate, as it produces numbers of pools that are a small multiple of a prime number (smaller than the prime itself).

***Light Weight***. Most theoretical pooling designs do not take into account the *weight* of the design, that is, the number of 1s in *M*. The weight is a critical feature for pooled experiments. Heavy weight means that each specimen is sampled more times, which consumes more material. In theory, of course, we can take a smaller amount of each specimen to mitigate this problem. But in reality, conventional liquid handling robots cannot accurately aspirate and dispense small quantities, limiting the possible weight. In addition, heavy weight designs are more laborious to implement and have higher overheads, contributing to the costs of the pooling procedure (robotic time, tips, or manual labor). In our experience and survey of the literature, tractable pooling designs rarely exceed a column weight of 10 in many cases.

***Modularity***. When the number of input specimens is large, a pooling experiment is typically divided into batches of distinct blocks of specimens. For each batch, the experimentalist essentially starts from scratch to construct the pools with the new subset of specimens. This approach is more robust because a sporadic failure of the liquid handling system affects only a limited number of specimens rather than the entire experiment. In addition, it limits the amount of liquid evaporation from the pools by minimizing the time it takes to finish constructing a pool. One desired property is that every batch will have a good performance and that the experimentalist will be able pool distinct batches together into a ‘mega-design’ without too much loss of performance. This will allow maximal flexibility in carrying out the multiple experiments. For instance, the experimenter can start with a conservative pilot experiment on one batch. Based on the results, she can decide whether to take a more aggressive approach and pool multiple batches together into a mega design or to retain the more conservative approach. Another advantage of modular designs stems from the fact that the constructed pools will typically be used many times in various conditions (e.g., using different bait proteins in yeast two-hybrid experiments), with varying numbers of positive items. A mega design can be sufficient in most cases, but decoding may fail when the number of positives is unusually high. However, when this happens one can redo the experiment using the underlying sub-designs, which have more smaller pools and will thus successfully identify the positives.

***Average performance***. Substantial parts of the theory of group testing address the worst-case performance of a pooling design, called the matrix disjunctness. With this property, it is very easy to derive the minimal number of positive items that can be decoded after pooling given a certain number of errors. In practice, however, the worst-case performance has low utility for experimentalists. What we really need is a design that performs well on average and to know the expected rather than worst case performance of the design.

***Robustness to Noise***. Much of classical group testing theory deals with the case of perfect measurements; i.e., we observe *y* = *Mx*. In practice, almost all measurements performed by experimentalists are noisy. We therefore need pooling designs and reconstruction algorithms which provide good average performance in a noisy setting. Up to now, these last two points have only been adressed by simulation [23].

**Reed-Solomon based pooling designs**. We have found a pooling design based on Reed-Solomon (RS) error correcting codes that dramatically reduces the number of pools compared to the pooling designs widely used in biological experiments, while addressing the practical properties mentioned above. In coding theory, a *code* is a set of vectors that differ pairwise in a large number of places. RS codes are *linear* codes, based on polynomials over *finite fields*.

Designs based on RS codes have been used in biological group testing since the 1960’s, when they were proposed by Kautz and Singleton [14]. However, existing work has not addressed the challenges above; the contribution of this work is showing how to use RS designs to address the practical needs of modern biology.

Below, we describe the construction of a RS design from the practitioner’s perspective; the supplementary material contains a more theoretical approach. We build the matrix *M* by stacking smaller submatrices, called *layers* on top of each other. The basic construction starts by selecting three integers: *q*, the number of pools in each layer, *m*, the number of layers, and *k*, the maximum degree of the polynomials we construct. We refer to such codes as [*m*, *k*, *m* − *k* + 1]*_q_*-RS codes. There are three constraints on these parameters, imposed by the theory. First, the number of specimens, *n*, must be at most *q_k_*; in practice, we will choose *k* to be the smallest integer so that this is true. Second, *q* must be either a prime or a prime power (e.g., 7, or 2^4^ = 16). For practical reasons discussed later, we are interested specifically in the case when *q* = 16. Third, both *k* and *m* must be smaller or equal to *q*. The pooling matrix *M* has *t* = *qm* rows (pools), and is divided into *m* submatrices (layers) of equal size *q* × *n*. We will number the layers 0, 1,…, *m* − 1, and we will number the rows within each layer 0, 1,…, *q* − 1.

Next, we represent each specimen number as a polynomial of degree *k*−1. We order the polynomials lexicographically. For example, if *k* = 3, and *q* = 16, then the first specimen is represented by the polynomial *P*_0_(*U*) = 0*U*^2^ + 0*U* + 0 = 0, the second specimen is represented by the polynomial *P*_1_(*U*) = 0*U*^2^ + 0*U* + 1 = 1, the 275th specimen is represented by *P*_275_(*U*) = *U*^2^ + *U* + 3, and the 276th by *P*_276_(*U*) = *U*^2^ + *U* + 4. Notice that this polynomial representation depends on *q*: if *q* = 7, then *P*_275_(*U*)=5*U*^2^ + 4*U* +2.

Each specimen will be assigned to one pool in each layer. To find the pooling assignment of the *i*th specimen in the *j*th layer, we will evaluate the polynomial *P_i_* at position *j*, *P_i_*(*j*), over the finite field of size *q*, denoted 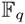. For example, if *q* = 7, *i* = 275, and *j* = 2, then *P*_275_(2) = 5 · 2^2^ + 4 · 2 + 3 mod 7 = 3.

Now, the *i* column in the *j* layer is a binary vector of length *q*, whose *ℓ*-th position is 1 if *P_i_*(*j*)= *ℓ* and 0 otherwise. Thus the *i*th sample will be assigned to the *ℓ*th pool in the *j*th layer. For example, if *q* = 7, in layer 2, the 275th specimen will be assigned to the pool labeled 3 in that layer.

We note that when *q* is a prime number, the addition and multiplication operations to evaluate the polynomial are simply performed modulo *q*. However, when *q* is a prime power, the arithmetic of the finite field is more involved—for example, over 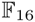, we have 4^2^ = 3. We have included in the supplementary material an introduction to finite fields and arithmetic tables for 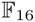. In this example, if *q* = 16, then we consider *P*_275_(2) = 2^2^ + 2 + 3 over 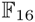, which turns out to be 5. Thus, the 275th specimen will be assigned to pool 5 in layer 2.

The process is illustrated in Figure 1, with *q* = 16, *m* = 6, and *n* = 288. We will return to this example throughout the paper to illustrate our ideas.

**Fig. 1.**
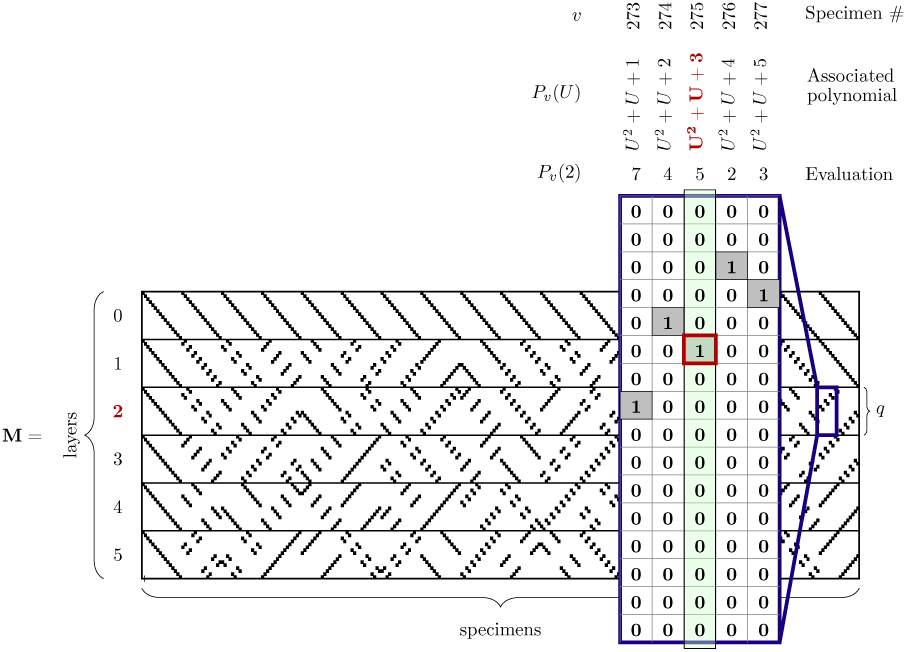
Construction of the Reed-Solomon pooling matrix with *q* = 16 and *m* =6, and *n* = 288. The columns corresponds to specimens and are indexed by polynomials over 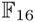, and the rows are grouped into 6 layers. We highlight the assignment of the specimen *v* = 275 in layer 2. As described in the text, we consider *P*_275_ = *U*^2^ + *U* +3, and evaluate *P*_275_(2) = 5.

**Fig. 2.**
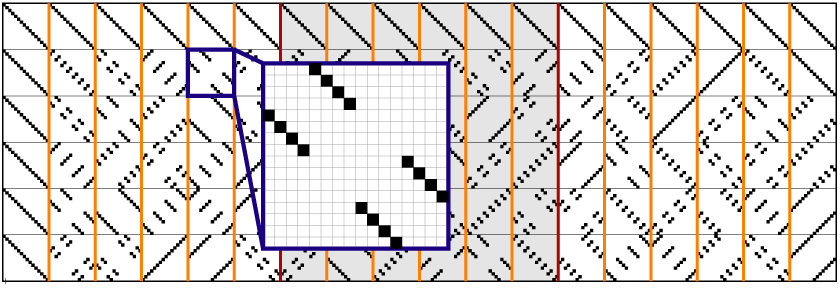
Returning to the example of Figure 1, we see that the resulting design matrix is *balanced*. In each 16 × 16 block, there is exactly one 1 in each row and each column. This implies that (a) each of the 288 specimens participates in exactly 6 pools, and (b) each pool contains exactly 288/16 = 18 specimens. Further, it implies that these properties hold modularly: the design given by the shaded submatrix has 96 specimens, each of which participates in exactly 6 pools, so that each pool contains exactly 6 of these specimens.

Shifted Transversal Designs [22] are special cases of RS codes, when *q* is limited to a prime rather than a prime power. We describe the correspondence in the supplementary material. We will show that RS codes possess all the qualities of STD, while providing more flexibility: contrary to STD, RS codes are instrinsically well–adapted to typical laboratory automation hardware.

## Structural Properties of Reed Solomon pooling designs

From the constructions of RS designs, we immediately see that several of our practical desires are met.

First, the parameter space is very rich, and in particular this allows for designs which fully utilize 96-well plates. As in the example above, choosing *m* = 6 and *q* = 16 yields precisely 96 wells. Further, as the specimens are also stored in 96-well plates, it is convenient for pooling designs to work with *n* specimens, where *n* is a multiple of 96. Again, the example above (where *n* = 288 = 3 · 96) demonstrates that this is easily obtainable. Further (as can be seen in Figure 1), these designs have a lot of structure, resulting in more efficient pooling as well as error checking of the pooling procedure by visual inspection.

Second, these designs enjoy excellent regularity and weight. Each specimen is pooled into exactly one pool in each layer; thus, each specimen participates in *m* pools total. It is also true that each pool contains exactly *n/q* specimens, as long as *n* is a multiple of *q*.

This fact implies very strong regularity properties: not only are the designs light and well-balanced, but these properties hold in a modular fashion. As described above, it is often useful to be able to pool different sets of specimens at different times. With RS designs, as long as the specimens are pooled in multiples of *q*, the scientist enjoys light and well-balanced designs in whatever sizes are convenient.

**Disjunctness and recovery**. In addition to convenient structure, the pooling design should allow for efficient recovery of the pattern of positive specimens, given the pooled data. For this, we introduce the notion of *disjunctness*. A pooling design is *d*-disjunct if for every set Λ of *d* specimens, and every specimen *i* not in that set, there is some pool *j* that includes *i* but doesn’t include any item in Λ. The resulting condition on the matrix *M* is illustrated in Figure 3. If *M* has this property, then we can quickly find the correct set of positives from the pooled results, assuming this set is not larger than *d*. We simply assert that an item *i* is positive if all of its pools were positive. If the item *i* was *not* positive, and the true set of positives was Λ, then the disjunctness condition precisely states that *i* will land in some pool that does not test positive. Disjunctness is discussed in more detail in the supplementary material.

**Fig. 3.**
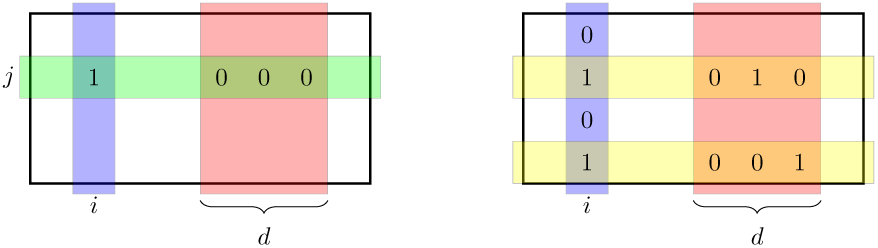
A pooling design is *d*-disjunct if for all Λ of size *d* and all *i*, there is a row *j* as in the lefthand diagram. In contrast, the matrix on the right is *not d*-disjunct.

For pooling designs arising from codes, like RS designs, it is easy to obtain a bound on disjunctness. The RS design defined above has disjunctness at least *d* = ⎿(*m* − 1)*/k*⏌. In our running example from Figure 1, this comes out to *d* =2. We will see in a later section (Proposition 2) why this is the case. In terms of the number of specimens *n*, the above bound guarantees that with *t* pools we can achieve disjunctness of a least 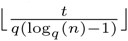. This bound gives a trade-off between recovery guarantees (the maximum number of defectives recovered, *d*) and budget (assumed to be proportional to the number of pools *t*). Moreover, the RS-based design described above is *incremental* - that is, an experimentalist can start with a small number of layers *m*, giving a small pooling matrix *M* with a certain disjunctness value, and gradually increase *m* by adding additional layers to the pooling matrix, hence increasing the disjuncness. Adding an additional layer *j* is easily achieved by evaluating the polynomials *p_i_* over the field 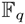 at the element 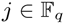.

## Modularity

As discussed in the introduction, in many applications it is convenient to pool samples in batches, so that these batches may either be tested independently, or mixed with others and tested jointly. More specifically, given two groups of *c* specimens each, one may pool each group separately (using two different pooling designs) into *p* pools each; the resulting design has 2*c* specimens pooled into 2*p* pools. Alternatively, one may combine or “superpose” the designs, resulting in 2*c* specimens pooled into *p* pools. The resulting situation for measurement matrices is shown in Figure 4.

**Fig. 4.**
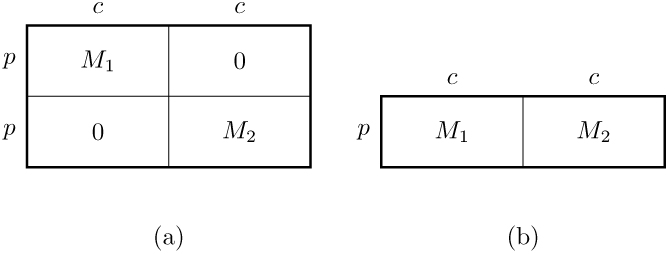
Different ways of combining two pooling designs given by matrices *M*_1_ and *M*_2_. In (a), each group of specimens is pooled separately, resulting in 2*p* pools. In (b), the pooling designs are combined, resulting in p pools. A design *M* consisting of submatrices *M*_1_,*M*_2_ has good *modularity* if the pooling matrix (a) has strictly better disjunctness than the matrix (b).

There are trade-offs between these two strategies: in the first, there are more tests but there are fewer specimens in each pool, potentially yielding more accurate assay results. In the second, there are fewer tests, but a possibility of less accuracy. One practical goal in pooling designs is to make this trade-off as beneficial as possible. That is, we would like to partition the columns of the pooling matrix *M* into submatrices *M_i_* so that each submatrix *M_i_* has good disjunctness. Trivially, each *M_i_* will be at least as disjunct as *M* itself, but we seek designs where passing to a submatrix yields a strict improvement in disjunctness (and hence a strictly better trade-off between the accuracy and the number of pools). We refer to this property as the *modularity* of the design, and we discuss the mathematical particulars in more detail in the section on general linear codes.

Even without improving disjunctness, the smaller designs corresponding to submatrices *M_i_* are easier to decode because the *d* positives will be spread across these subdesigns, so that each experiment will have to deal with (and identify) fewer positives.

In practice, there are many reasons that one might wish to have the kind of flexibility that modularity entails. For example, suppose—as is often the case—that one does not know the true number of positives, *d*. Then one may use modularity to estimate *d* on the fly, as follows. Suppose that the pooling design is given by a modular matrix *M*, which decomposes into submatrices *M*_1_*,M*_2_,…,*M_b_*, where each submatrix *M_i_* assigns, say, 96 items to *p* pools. Before doing any tests, the biologist may pool her samples into *b* different 96-well plates, according to the *b* different pooling designs, each with very good disjunctness. When it is time to run tests, she may choose to start with a single plate; if her results are very good, she may think that she has over-estimated *d*. For her next two tests, she may choose to combine the second and third plates, which results in a design (as in Figure 4(b)), with worse disjunctness but fewer pools. If this still appears too pessimistic, she may continue in this way, testing more and more plates at a time until she hits the desired trade-off between disjunctness and the number of pools.

As another example, the Shifted Transversal Design has good modularity, and this has provided important practical benefits in an interactome mapping experiment [25]. In this work, subdesigns (“micro-pools”) were constructed once, and were then combined by superposition to obtain designs that were well–suited to various experimental formats: in pairs for the 1536–well format, in sixes for the 384–well format, and in twelves for the 96–well format.

RS designs (and, as is the trend in this work, designs from appropriate linear codes more generally) have good modularity properties. The basic idea is that each slab of *q* columns (demarcated by the orange lines in Figure 2) has similar disjunctness properties. This follows from the fact that the sets of polynomials they correspond to look similar: as far as disjunctness is concerned, there is not much difference between the set {*U*^2^ + *U*, *U*^2^ + *U* +1,…,*U*^2^ + *U* + 15}and the set of constant polynomials {0, 1,…, 15}. Similarly, at a larger scale, there is not that much difference between the set of all polynomials of the form *U*^2^ + *aU* + *b* and the set of all polynomials of the form 2*U*^2^ + *aU* + *b*.

Because of the nature of these symmetries, the modularity naturally occurs for sets of columns of size *q*, *q*^2^, *q*^3^, and so on. For example, when *q* = 16, *m* =6, and *k* = 3, we may support up to *n* = *q^k^* = 16^3^ = 4096 items with disjunctness 2. These items can be divided into subsets of size 16^2^ = 256 with a guarantee of disjunctess 5. However, as discussed above, it is often convenient if the sizes of the subsets of specimens are multiples of 96. For illustration purposes, consider a slightly larger version of our running example, where *q*, *m*, and *k* are as above, but where *n* = 576 =6 · 96. If we split up the specimens into six batches of size 96 in the natural way: the first 96 columns, then the next 96 and so on, most of the blocks of size 96 do not interfere with the blocks of size 256 (or do not overlap the 256-block boundaries). Thus, most batches of 96 specimens obtained this way inherit the improved disjunctness of the blocks of size 256. This same argument works for any multiple of 96.

## Average disjunctness

Whether we exploit the modularity of the RS design or not, the entire design of our running example has disjunctness of only two which seems rather bleak. Unfortunately, with only 96 pools and a large number of samples, it is not possible to obtain much better worst-case guarantees. However, in practice, these guarantees are often overly pessimistic. Thus, it is natural to ask about the performance of these designs against *most* patterns of positives. We seek a statement of the form, “for *most* patterns of defectives, the group testing matrix is effectively *d*-disjunct.”

To this end, we define a weaker notion of disjunctness, which we call the *average disjunctness*, which measures how many defectives can be reliably identified in a “typical” rather than worst-case situation. We say that a set Λ of specimens is *bad* if it causes a situation as in the right-hand side of Figure 3. That is, Λ is bad if there is some specimen *i* so that *i* coincides with at least one member of Λ each time it is pooled. Thus, a design is *d*-disjunct if all sets Λ of size *d* are not bad. We relax this in a natural way, and say that a pooling design is (*δ*, *d*)*-average-disjunct*, if at most a *δ*-fraction of the sets of specimens Λ of size *d* are bad. (When *δ* is not mentioned, we take it to be an appropriate small constant, like 0.01).

Thus, if the positive specimens are randomly distributed, with probability 1−*δ*, the set of positive specimens that appears is not bad. Thus, by the same reasoning about the (worst-case) disjunctness, we conclude that (*δ*, *d*)-average-disjunctness allows efficient identification of *d* positives chosen uniformly at random with probability at least 1 − *δ*. Of course, it is unreasonable to assume in practice that the positive specimens will be uniformly distributed. However, the above reasoning works for any distribution which is “close” to the uniform one in an appropriate sense; e.g., physical specimens are assigned column identities at random.

A matrix may be *d*-average-disjunct without being *d*-disjunct, and we are interested in quantifying and exploiting this gap. We do so both in practice and in theory. In the supplementary material, we show how to efficiently compute simple, provable bounds on the values of (*δ*, *d*) so that RS designs are (*δ*, *d*)-average disjunct. We complement our theoretical results with empirical ones, which indicate that our bounds are quite accurate for small *d*, and lose accuracy as *d* grows.

Our results, both theoretical and empirical, are displayed in Figure 5. We compared the empirical performance of the RS pooling design to two randomized designs. For the first random design, we chose *n* random columns from an RS design with the full *q^k^* possible columns; this comparison is meant to illustrate that our modular column selection is not only convenient for practical considerations, it also actually yields better performance. For the second random design, we chose a random *m* · *q* × *n* matrix with *m* ones per column. Theoretically, random matrices offer the best asymptotic guarantees, and so the fact that RS designs outperform it for reasonably sized values of *n* is striking.

**Fig. 5.**
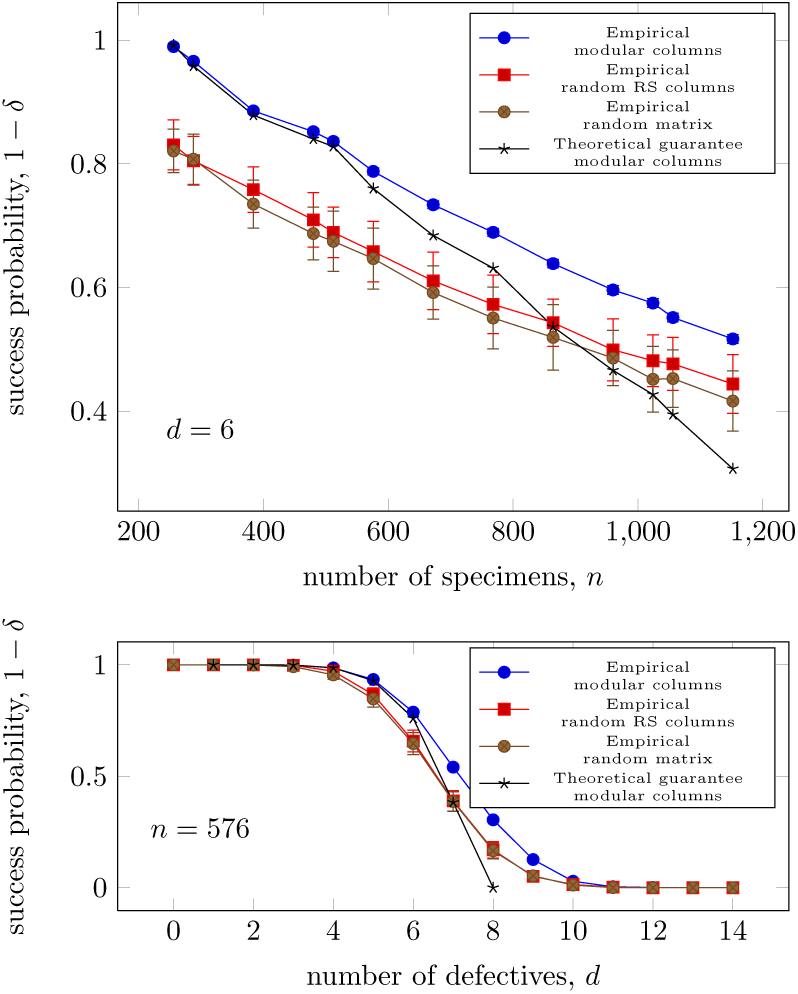
Empirical and theoretical bounds on (*δ*, *d*)-average disjunctness for RS designs. In both charts, the empirical performance of the RS pooling design is compared to its theoretical performance, as well as the empirical performance of two randomized pooling designs. For the RS matrices, empirical performance was judged using 5000 random trials. For the random matrices, empirical performance was judged from 100 trials each on 100 random matrices. The theoretical bounds were obtained through a recursive formula described in the supplementary material. The top chart traces the empirical and provable success probability 1 − *δ* of a RS design with *m* = 6 and *d* = 6, as the number of specimens *n* grows, and compares this to the failure probability of random designs with the same column weight and number of tests. The bottom chart fixes *n* = 576 = 6 · 96, and varies *d*. For all RS designs, the worst-case disjunctness is two.

These results are much more hopeful than the worst-case bounds of *d* = 2. For *n* = 576, for example, up to 7 or 8 positives may be accurately identified with reasonable probability. Further, we see that RS designs perform much better than comparable randomly generated designs.

## Experimental validation of RS-based pooling

We tested the RS design with an experiment that combined 480 samples into 96 pools (*m* =6, *q* = 16). For this experiment, we used a toy problem in which individual samples were represented by a single well of 150 *µ*1 of deionized water and positive samples of water were spiked with blue dye (100X dilution). The aim of the experiment was to find the positive (blue dye) specimens by inspecting the pools. Pooling was carried out with a Tecan Evo robot using a four-tip liquid-handling arm (LiHa). To acquire the data, we quantified the absorbance ratio (470nm/630nm) of each pool using a plate reader. Then, we used an out-of-the-box conventional compressed sensing decoder (GPSR) [19] to decode the results.

Despite the molecular simplicity, our experimental design captures some of the challenges inherent to pooling experiments. First, it tests the actual execution in a laboratory setting. Second, this simple experiment captures the three main challenges of pooling experiments: pipetting, dilution, and measurement noise. Pipetting noise refers to the inability of the experimentalist to aspirate and dispense the exact volume of the input specimen due to human error. These imperfections introduce noise to the pooling matrix that can distort the signal. Based on our empirical results, the coefficient of variation of the pipetting steps was 5–10%. Dilution noise occurs when the signal from the positive specimen is diminished as the size of the pool increases. In our toy experiment, each pool contained 30 specimens, diluting the blue dye signal by the same factor. Measurement noise in this example would result from the absorbance scan to measure the blue dye of the constructed pools. In this case, the measurement noise mainly originated from the number of photons of each scanned pool.

Despite these imperfections, we were able to accurately decode the pooled results and identify the original specimens when three were positive. We repeated this experiment with ten positive specimens and were able to correctly decode eight of the original specimens with one false positive and one false negative. This is not surprising as simulations shown in Figure 5 exhibit imperfect decoding for similar conditions.

Encouraged by these results, we also tested our pooling design to find rare genetic variations using targeted high throughput DNA sequencing. Targeted sequencing is a labor intensive and costly procedure that processes DNA in multiple steps to enrich for specific genomic regions; instead, we used a pooling design. We pooled equimolar amounts of 96 DNA samples from the HapMap Project into 25 pools using an RS code with *m* =5 and *q* =5 (also a Shifted Transversal Design). The 25 pools were barcoded and enriched by hybrid selection using Agilent SureSelect baits that enriched for genes found in the Framingham Heart Study (≈ 790 kb of genomic material). The pools were further compressed into five mega pools and sequenced on five lanes on the Illumina HiSeq (2 × 80 bp) to 1000 times the average coverage per pool.

To test our approach, we examined 75 of our samples that were also sequenced as part of the 1000 Genomes Project [4]. More than 97% of the target region was covered by at least one sequencing read. After filtering positions with low quality data (see supplementary material), we were left with 92% of the bases in the original design. Next, we exploited the structure of the RS code to estimate the number of carriers in our design before decoding and attributing the mutation to individual specimens. This procedure relies on the observation that each specimen participates in exactly one pool per layer. By inspecting the number of variant reads across pools in the same layer, it is possible to estimate the number of carriers. Since our design had five layers, we repeated this procedure five times and averaged the results to get more accurate estimates. We compared the number of carriers between our estimator using the pooling data and an estimator using the individual-level results of the 75 specimens in the 1000 Genomes data. Figure 6 shows excellent consistency (*R*^2^ = 0.91) between the two estimators (perfect consistency cannot be observed since we have individual level data for only 75 out of 96 samples).

**Fig. 6.**
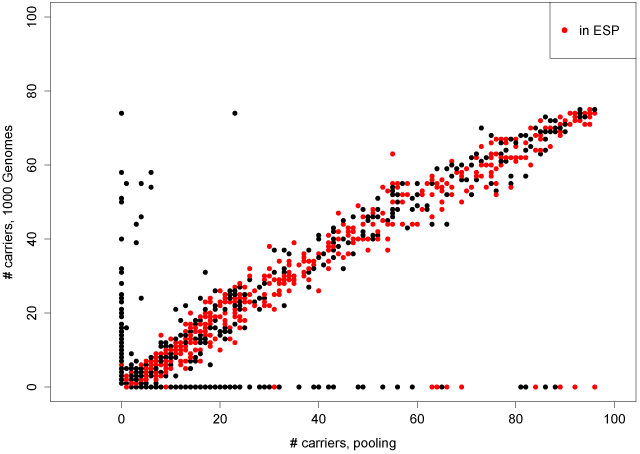
Comparison of carrier rates from 75 HapMap samples between 1000 Genomes and our pooled samples. Further comparison to the Exome Sequencing Project (variable sites are red) suggests that misdetections in our pooled samples are false positives in 1000 Genomes.

Our results also found good decoding capabilities for rare variants. We decoded the pooled data for variants where our pooled estimator indicated five or fewer carriers (translating to a minor allele frequency of 2%). We used a series of decoders that took into account the sequencing noise at each position and the distribution of reads with the variant allele. These decoders ran on each nucleotide separately and used the pooling matrix to find which specimens actually carry the rare allele. Our results indicate a specificity of > 99.99% and sensitivity of about 85% compared to the 1000 Genomes data, averaged across all positions. We wondered if the false negatives represent a true decoding miss or a sequencing error in the 1000 Genomes project due to the low coverage of these genomes. We turned to the National Heart, Lung, and Blood Institute (NHLBI) Exome Sequencing Project (ESP) [8], which contains genomic data for more than 5000 individuals. The expectation is that sequencing errors in the 1000 Genomes data will not appear as variable sites in ESP. Indeed, when we inspected missed positions in our results only 26% of them were variable in ESP. We also inspected positions where our decoder did find a rare variant signal in the pooled data. Close to 38% of those positions were also variable in ESP. This suggests that some of the imperfect sensitivity can be attributed to sequencing errors in the reference data rather than a problem with the decoder.

## Tools for Experimentalists

We created a free web tool to design and guide experiments (http://pooling.teamerlich.org). A user uploads a sample map spreadsheet and enters the number of specimens and volume constraints. Then, she can select her desired pooling design (for completeness and education purposes, our tool can generate several pooling designs that are described in the group testing literature in addition to the standard RS code design (*q* = 16, *m* =6). Based on user input, the tool checks that the specimens have sufficient volume to be pooled and that the pooling plate can hold the accumulated volume from the input material without spillover. Then, the tool generates a design specific matrix and pooling instructions.

The pooling instructions are compatible with a liquid-handling robot or manual hardware such as iPipet [26]. The latter is a semi-automated open-source tool that we developed to convert a regular iPad (and most tablets) into a sample-tracking device. Users simply place the sample plate and pooling plate on the tablet, and iPipet illuminates the wells to be aspirated or dispensed. Based on our tests, a trained experimentalist can complete an RS-code based pooling experiment of ≈ 500 input specimens within a few hours using iPipet. With this tool and the theory outlined in this paper, we hope to reduce the technical barriers for experimentalists to conduct pooled experiments and to provide guidance based on rigorous theoretical analysis.

## ACKNOWLEDGMENTS

The authors would like to thank the Institute for Mathematics and its Applications for facilitating our initial collaboration. Y.E. holds a Career Award at the Scientific Interface from the Burroughs Wellcome Fund. This study was supported by a gift from Andria and Paul Heafy and by NIH grant R21HG006167 and National Science Foundation grants CCF-1161196, CCF 1161233, and CCF 0910765. The order of the authors follows the convention in Mathematics and is alphabetical. All authors contributed equally to the paper.

## Appendix

**Table 1.** Summary of biological assays that use pooling designs.

| Aim | items ($x$) | assay ($y$) | Pooling design | Reference |
| --- | --- | --- | --- | --- |
| Finding sequencing tag sites | Yeast artificial chromosome library (YAC) | PCR amplification | k-set packing design | [2] |
| Gene expression | Samples of *Arabidopsis thaliana* | Expression array | Expander | [13] |
| Genome assembly | BACs of *Barley* | High throughput sequencing | STD | [17] |
| Interactome mapping | Yeast strains expressing an OR-Feome as Y2H prey constructs | Yeast two hybrid | STD | [25] |
| Interactome mapping | Protein domains | Protein microarray | logarithmic pattern | [12] |
| Interactome mapping | Yeast strains expressing an OR-Feome as Y2H prey constructs | Yeast two hybrid | logarithmic pattern | [11] |
| Mutant screens for drug resistance | Yeast library with systematic deletions | growth assay | logarithmic pattern | [12] |
| Validation of RNAi libraries | RNAi inserts in bacterial colonies | high throughput sequencing | CRT | [7] |
| Fitness profiling | Fission yeast deletion strains | high throughput sequencing | STD | [10] |
| Discovery of synergistic drug combinations | Chemical compounds | Fluorescence-based assay | STD | [21] |

**Mathematical background and definitions**. We begin with some notation. For any integer *n* ≥ 1, let [*n*] denote the set {1,…,*n*}. Given an *n*-dimensional vector x, we will denote its *i*th component by *x_i_*, *i* ∈ [*n*]. The entry of a matrix *M* in the *i*th row and *j*th column will be denoted by *M_i_*,*_j_*.

For any set *X* of numbers and any positive integer *n*, let *X^n^* denote the set of all *n*-dimensional vectors v =(*v*_1_*,v*_2_, ···*, v_n_*) such that each *v_i_*, *i* ∈ [*n*], is a member of *X*. For example, ℝ*^n^* is the set of all *n*-dimensional real vectors, and 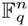 is the set of all *n*-dimensional vectors over the finite field 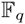. (We briefly define finite fields below.)

**Finite fields**. Informally, a *field* is a set of elements such that one can perform addition, multiplication, subtraction, and division on these elements without obtaining results outside the set. For example, the set of real numbers is a field. These properties of a field endow the real numbers with the familiar set of arithmetic properties. When the field is finite, the structure becomes quite different. In order for the results of the four operations to always stay within the set, both the set size and the meanings of addition, subtraction, division, and multiplication have to have a specific structure. For example, a finite field can only have size *q*, where *q* is a prime power. We shall use 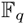 to denote the finite field on *q* elements, as it is a fundamental result in algebra that all finite fields of the same size are isomorphic—that is, they have the same structure. In every finite field 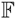, there is a designated element denoted by 0 and another denoted by 1. Every element 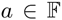 has an *additive inverse b* ∈ *F* such that *a* + *b* =0. Every element 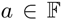 with *a* ≠ 0 has a *multiplicative inverse* 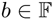 such that *a*·*b* = 1. Sometimes, we denote the additive inverse of *a* by −*a*, and the multiplicative inverse of *a* by *a*^−1^. Then, subtraction and division are defined to be equivalent to addition by the additive inverse and multiplication by multiplicative inverse. This way, only addition and multiplication need to be formally defined for a finite field 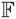. Addition and multiplication in a finite field can be computed using an addition and a multiplication table, as shown in an example below. A full discussion of finite fields, their operations, their addition and multiplication tables, their properties, and the motivations for having them will require a full textbook on abstract algebra (e.g., [15]). Within the scope of this supplementary material, we can only present a sketch of finite field properties we need to define our pooling designs. We next give examples of specific finite fields.

**The finite field** 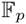 **where *p* is a prime number**. Finite fields 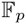 where *p* is a prime are called *prime fields*. The set of elements are 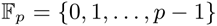, where the addition (let us denote it by +) is normal addition modulo *p* (i.e. we add the numbers as integers and then take the remainder when we divide the sum by *p*) and multiplication (let us denote it by ·) is normal multiplication modulo *p* (i.e. we multiply the numbers as integers and then take the remainder when we divide the product by *p*). Subtraction and division are correponding inverses of addition and multiplication. It can be shown that in this case the inverses are well-defined (except for division by 0).

We illustrate prime fields with *p* = 13. Tables 3 and 4 give the tables of addition and multiplication of two elements from 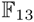 respectively.

**The fields** 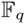 **for *q* power of 2**. For designing practical group testing matrices, we often work with fields 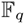 where *q* is a power of 2. In this case the arithmetic operations in 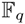 are a bit more complicated than the prime field case. For example, when *q* = 2^4^ = 16, Tables 5 and 6 present the tables for addition and multiplication of two elements from 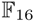 respectively.

**Table 2.** Specific notation for quantities of interest.

|  |  |
| --- | --- |
| $m$ | number of polynomial evaluation points |
| $t$ | number of pools or tests |
| $q$ | field size |
| $k$ | maximum degree of polynomials in RS code |
| $n$ | total number of polynomials in RS code |
| $n$ | number of items or specimens |
| $m$ | number of layers in RS matrix |

**Table 3.** Table for addition over 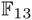

| + | 0 | 1 | 2 | 3 | 4 | 5 | 6 | 7 | 8 | 9 | 10 | 11 | 12 |
| --- | --- | --- | --- | --- | --- | --- | --- | --- | --- | --- | --- | --- | --- |
| 0 | 0 | 1 | 2 | 3 | 4 | 5 | 6 | 7 | 8 | 9 | 10 | 11 | 12 |
| 1 | 1 | 2 | 3 | 4 | 5 | 6 | 7 | 8 | 9 | 10 | 11 | 12 | 0 |
| 2 | 2 | 3 | 4 | 5 | 6 | 7 | 8 | 9 | 10 | 11 | 12 | 0 | 1 |
| 3 | 3 | 4 | 5 | 6 | 7 | 8 | 9 | 10 | 11 | 12 | 0 | 1 | 2 |
| 4 | 4 | 5 | 6 | 7 | 8 | 9 | 10 | 11 | 12 | 0 | 1 | 2 | 3 |
| 5 | 5 | 6 | 7 | 8 | 9 | 10 | 11 | 12 | 0 | 1 | 2 | 3 | 4 |
| 6 | 6 | 7 | 8 | 9 | 10 | 11 | 12 | 0 | 1 | 2 | 3 | 4 | 5 |
| 7 | 7 | 8 | 9 | 10 | 11 | 12 | 0 | 1 | 2 | 3 | 4 | 5 | 6 |
| 8 | 8 | 9 | 10 | 11 | 12 | 0 | 1 | 2 | 3 | 4 | 5 | 6 | 7 |
| 9 | 9 | 10 | 11 | 12 | 0 | 1 | 2 | 3 | 4 | 5 | 6 | 7 | 8 |
| 10 | 10 | 11 | 12 | 0 | 1 | 2 | 3 | 4 | 5 | 6 | 7 | 8 | 9 |
| 11 | 11 | 12 | 0 | 1 | 2 | 3 | 4 | 5 | 6 | 7 | 8 | 9 | 10 |
| 12 | 12 | 0 | 1 | 2 | 3 | 4 | 5 | 6 | 7 | 8 | 9 | 10 | 11 |

**Table 4.** Table for multiplication over 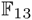

| · | 0 | 1 | 2 | 3 | 4 | 5 | 6 | 7 | 8 | 9 | 10 | 11 | 12 |
| --- | --- | --- | --- | --- | --- | --- | --- | --- | --- | --- | --- | --- | --- |
| 0 | 0 | 0 | 0 | 0 | 0 | 0 | 0 | 0 | 0 | 0 | 0 | 0 | 0 |
| 1 | 0 | 1 | 2 | 3 | 4 | 5 | 6 | 7 | 8 | 9 | 10 | 11 | 12 |
| 2 | 0 | 2 | 4 | 6 | 8 | 10 | 12 | 1 | 3 | 5 | 7 | 9 | 11 |
| 3 | 0 | 3 | 6 | 9 | 12 | 2 | 5 | 8 | 11 | 1 | 4 | 7 | 10 |
| 4 | 0 | 4 | 8 | 12 | 3 | 7 | 11 | 2 | 6 | 10 | 1 | 5 | 9 |
| 5 | 0 | 5 | 10 | 2 | 7 | 12 | 4 | 9 | 1 | 6 | 11 | 3 | 8 |
| 6 | 0 | 6 | 12 | 5 | 11 | 4 | 10 | 3 | 9 | 2 | 8 | 1 | 7 |
| 7 | 0 | 7 | 1 | 8 | 2 | 9 | 3 | 10 | 4 | 11 | 5 | 12 | 6 |
| 8 | 0 | 8 | 3 | 11 | 6 | 1 | 9 | 4 | 12 | 7 | 2 | 10 | 5 |
| 9 | 0 | 9 | 5 | 1 | 10 | 6 | 2 | 11 | 7 | 3 | 12 | 8 | 4 |
| 10 | 0 | 10 | 7 | 4 | 1 | 11 | 8 | 5 | 2 | 12 | 9 | 6 | 3 |
| 11 | 0 | 11 | 9 | 7 | 5 | 3 | 1 | 12 | 10 | 8 | 6 | 4 | 2 |
| 12 | 0 | 12 | 11 | 10 | 9 | 8 | 7 | 6 | 5 | 4 | 3 | 2 | 1 |

**Table 5.** Table for addition over 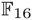

| + | 0 | 1 | 2 | 3 | 4 | 5 | 6 | 7 | 8 | 9 | 10 | 11 | 12 | 13 | 14 | 15 |
| --- | --- | --- | --- | --- | --- | --- | --- | --- | --- | --- | --- | --- | --- | --- | --- | --- |
| 0 | 0 | 1 | 2 | 3 | 4 | 5 | 6 | 7 | 8 | 9 | 10 | 11 | 12 | 13 | 14 | 15 |
| 1 | 1 | 0 | 3 | 2 | 5 | 4 | 7 | 6 | 9 | 8 | 11 | 10 | 13 | 12 | 15 | 14 |
| 2 | 2 | 3 | 0 | 1 | 6 | 7 | 4 | 5 | 10 | 11 | 8 | 9 | 14 | 15 | 12 | 13 |
| 3 | 3 | 2 | 1 | 0 | 7 | 6 | 5 | 4 | 11 | 10 | 9 | 8 | 15 | 14 | 13 | 12 |
| 4 | 4 | 5 | 6 | 7 | 0 | 1 | 2 | 3 | 12 | 13 | 14 | 15 | 8 | 9 | 10 | 11 |
| 5 | 5 | 4 | 7 | 6 | 1 | 0 | 3 | 2 | 13 | 12 | 15 | 14 | 9 | 8 | 11 | 10 |
| 6 | 6 | 7 | 4 | 5 | 2 | 3 | 0 | 1 | 14 | 15 | 12 | 13 | 10 | 11 | 8 | 9 |
| 7 | 7 | 6 | 5 | 4 | 3 | 2 | 1 | 0 | 15 | 14 | 13 | 12 | 11 | 10 | 9 | 8 |
| 8 | 8 | 9 | 10 | 11 | 12 | 13 | 14 | 15 | 0 | 1 | 2 | 3 | 4 | 5 | 6 | 7 |
| 9 | 9 | 8 | 11 | 10 | 13 | 12 | 15 | 14 | 1 | 0 | 3 | 2 | 5 | 4 | 7 | 6 |
| 10 | 10 | 11 | 8 | 9 | 14 | 15 | 12 | 13 | 2 | 3 | 0 | 1 | 6 | 7 | 4 | 5 |
| 11 | 11 | 10 | 9 | 8 | 15 | 14 | 13 | 12 | 3 | 2 | 1 | 0 | 7 | 6 | 5 | 4 |
| 12 | 12 | 13 | 14 | 15 | 8 | 9 | 10 | 11 | 4 | 5 | 6 | 7 | 0 | 1 | 2 | 3 |
| 13 | 13 | 12 | 15 | 14 | 9 | 8 | 11 | 10 | 5 | 4 | 7 | 6 | 1 | 0 | 3 | 2 |
| 14 | 14 | 15 | 12 | 13 | 10 | 11 | 8 | 9 | 6 | 7 | 4 | 5 | 2 | 3 | 0 | 1 |
| 15 | 15 | 14 | 13 | 12 | 11 | 10 | 9 | 8 | 7 | 6 | 5 | 4 | 3 | 2 | 1 | 0 |

**Table 6.** Table for multiplication over 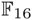

| · | 0 | 1 | 2 | 3 | 4 | 5 | 6 | 7 | 8 | 9 | 10 | 11 | 12 | 13 | 14 | 15 |
| --- | --- | --- | --- | --- | --- | --- | --- | --- | --- | --- | --- | --- | --- | --- | --- | --- |
| 0 | 0 | 0 | 0 | 0 | 0 | 0 | 0 | 0 | 0 | 0 | 0 | 0 | 0 | 0 | 0 | 0 |
| 1 | 0 | 1 | 2 | 3 | 4 | 5 | 6 | 7 | 8 | 9 | 10 | 11 | 12 | 13 | 14 | 15 |
| 2 | 0 | 2 | 4 | 6 | 8 | 10 | 12 | 14 | 3 | 1 | 7 | 5 | 11 | 9 | 15 | 13 |
| 3 | 0 | 3 | 6 | 5 | 12 | 15 | 10 | 9 | 11 | 8 | 13 | 14 | 7 | 4 | 1 | 2 |
| 4 | 0 | 4 | 8 | 12 | 3 | 7 | 11 | 15 | 6 | 2 | 14 | 10 | 5 | 1 | 13 | 9 |
| 5 | 0 | 5 | 10 | 15 | 7 | 2 | 13 | 8 | 14 | 11 | 4 | 1 | 9 | 12 | 3 | 6 |
| 6 | 0 | 6 | 12 | 10 | 11 | 13 | 7 | 1 | 5 | 3 | 9 | 15 | 14 | 8 | 2 | 4 |
| 7 | 0 | 7 | 14 | 9 | 15 | 8 | 1 | 6 | 13 | 10 | 3 | 4 | 2 | 5 | 12 | 11 |
| 8 | 0 | 8 | 3 | 11 | 6 | 14 | 5 | 13 | 12 | 4 | 15 | 7 | 10 | 2 | 9 | 1 |
| 9 | 0 | 9 | 1 | 8 | 2 | 11 | 3 | 10 | 4 | 13 | 5 | 12 | 6 | 15 | 7 | 14 |
| 10 | 0 | 10 | 7 | 13 | 14 | 4 | 9 | 3 | 15 | 5 | 8 | 2 | 1 | 11 | 6 | 12 |
| 11 | 0 | 11 | 5 | 14 | 10 | 1 | 15 | 4 | 7 | 12 | 2 | 9 | 13 | 6 | 8 | 3 |
| 12 | 0 | 12 | 11 | 7 | 5 | 9 | 14 | 2 | 10 | 6 | 1 | 13 | 15 | 3 | 4 | 8 |
| 13 | 0 | 13 | 9 | 4 | 1 | 12 | 8 | 5 | 2 | 15 | 11 | 6 | 3 | 14 | 10 | 7 |
| 14 | 0 | 14 | 15 | 1 | 13 | 3 | 2 | 12 | 9 | 7 | 6 | 8 | 4 | 10 | 11 | 5 |
| 15 | 0 | 15 | 13 | 2 | 9 | 6 | 4 | 11 | 1 | 14 | 12 | 3 | 8 | 7 | 5 | 10 |

**Linear codes**. Most of the results on Reed-Solomon codes extend to pooling designs generated from *linear codes*. We first sketch what we mean by linear codes for a general audience and then give more technical details.

Another way of viewing the construction of RS designs is as follows. First, we construct an intermediate *m* × *q^k^*^+1^ matrix 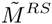 with entries in 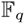. As before, the columns are indexed by polynomials *P* of degree at most *k*. The rows are indexed by the first *m* elements of 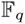. The entry in row *i* and column *P* contains (*i*). In order to obtain our design matrix *M* = *M^RS^*, we turn each row of 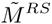 into a layer, replacing 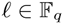 with a column that is 1 in the *ℓ*’th position and zero elsewhere. Finally, we take only the first *n* columns.

The columns of 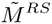 form a *Reed-Solomon code*, which is why we have called *M^RS^* an RS design. We may perform the same procedure on 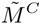, where the columns of 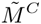 form any set 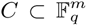, which we call a *code*. Reed-Solomon codes are *linear*, which means that the columns of *M^RS^* are closed under addition (over 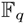), and we will be interested in codes *C* with this property. One parameter of interest of a code 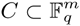 is the *distance*, that is, the maximum Hamming distance (number of differing symbols) between any two columns of *M^C^*. For example, the distance of Reed-Solomon codes is *m* − *k*, because any two polynomials *P* and *Q* of degree at most *k* can agree in at most *k* places. Any linear code may be described by an *s* × *m generator matrix*, and *s* is called the *dimension* of *C*. For example, the dimension of a Reed-Solomon code with degree *k* is *k* +1. If 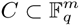 has distance Δ and dimension *s*, we say it is a [*m*, *s*, Δ]*_q_* code.

More formally, an [*m*, *k*, Δ]*_q_-linear-code* is a set *C* of *m*-dimensional vectors over the finite field 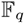 (that is, 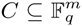), so that *C* forms a *k*-dimensional subspace of 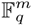.

Elements of *C* are called *codewords* of the code. Each codeword can be identified by a *message*, which is a vector 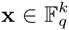. In particular, 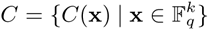 and thus |*C*| = *q^k^*. We can think of a message x as an index to access a code array *C*. Each member of the array is a vector in 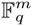 and there are *q^k^* members in the array. Because *C*(x) forms a subspace, the codewords *C*(x) are determined by a full rank *k* ×*m* matrix *G* over 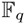, called the *generator matrix* of the linear code. More precisely, for every message 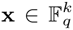, the corresponding codeword 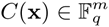 is defined by *C*(x)= x · *G*. Here, all arithmetic is carried out over the finite field 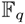.

The *weight* of a codeword is the number of nonzero components. The *Hamming distance* Δ(c, c^′^) between two codewords c ≠ c^′^ ∈ *C* is the number of coordinates *i* ∈ [*m*] such that 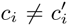. The *minimum distance*, or simply *distance*, of the code *C* is the minimum Hamming distance between two different codewords in *C*. For an [*m*, *k*, Δ]*_q_*-linear-code, Δ is the distance. Equivalently, because the code is linear (i.e., it can be defined with a generator matrix), it has minimum distance Δ if and only if the minimum weight of non-zero codewords is Δ.

### Definition 1

*A* coset *of an* [*m*, *k*, Δ]*_q_-linear-code C is a set of vectors of the form 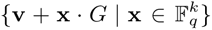*, *where* 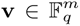 *is a fixed (but arbitrary) vector. The coset is often denoted by* v + *C*.

It can be shown that for two vectors **v**, 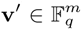, the cosets **v** + *C* and **v**′ + *C* are either identical or disjoint. And, it is well-known that the disjoint cosets of *C* form a partition of the vector space 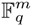.

### Definition 2

*Given a linear code C with generator matrix G*, *the* dual linear code *C*^⊥^ *of C is defined to be the set*

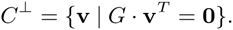

*In other words*, *the dual linear code consists of all vectors in* 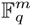 *which are orthogonal to all vectors in C*.

Clearly, Reed-Solomon codes are a special case of linear codes and we give a formal definiton of RS codes.

### Definition 3

(Reed-Solomon Code). *Let* 1 ≤ *k* ≤ *m* ≤ *q be positive integers such that q is a prime power. An* [*m*, *k*, *m* − *k* + 1]*_q_ Reed-Solomon (*RS*) code is defined as follows. Pick some set S of m distinct elements* 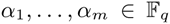 *called the* evaluation points*. Then*, *associate each message* 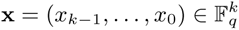 *with the univariate polynomial*

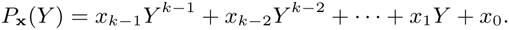

*Finally*, *the Reed-Solomon codeword corresponding to* x *is the vector obtained by evaluating P*_x_(*Y*) *at the n chosen points; i.e*.,

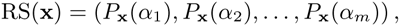
*and we define the Reed-Solomon code as*

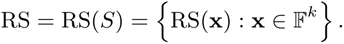

Given this definition, it is not too hard to see that the generator matrix corresponding to the above Reed-Solomon code is

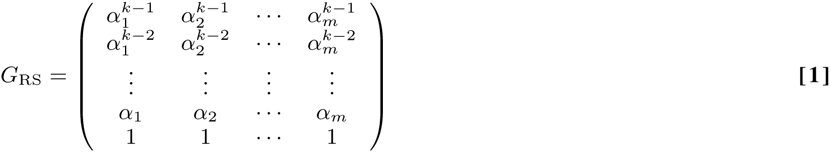

We will also consider subsets of Reed-Solomon codes, which correspond to picking a subset Ω of degree-(*k* − 1) polynomials (or equivalently a subset of messages 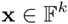). We refer to the subset of the Reed-Solomon code obtained by using polynomials Ω and evaluation points *S* as

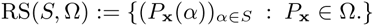

### Definition 4

(Binary matrix from linear code or a coset of a linear code). *Given an* [*m*, *k*, Δ]*_q_ linear code C*, *we define an* (*mq*) × *q^k^ matrix M^C^ as follows. We associate each of the mq rows with a pair* 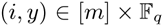, *and each of the q _k_ columns with a message 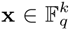*x/>*. Then we define the entry of M^C^ in row* (*i*, *y*) *and column* x *by*

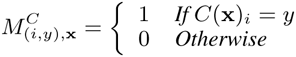

*Let* 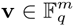 *be an arbitrary vector. We define the binary matrix M*^v+^*^C^ in the same way:*

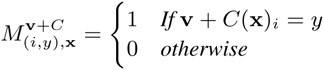

We conclude this section by using the definitions of a Reed-Solomon code and its associated binary matrix to show that the pooling design given by the binary matrix of a special case of a Reed-Solomon code is equivalent to the shifted transversal design (STD), a design used in a number of biological applications.

### Definition 5

*Let n be the number of desired columns*, *choose a prime number p with p* < *n*, *set k equal to the smallest integer κ such that p^κ^* ≥ *n*, *and choose the desired number of layers m* ≤ *p*., *The shifted transversal design STD*(*n*, *p*, *m*) *construction from [22] is defined as the concatenation of m p* × *n Boolean matrices*, *M*^(^*^ℓ^*^)^, *for ℓ* = 0,…,*m* − 1*; i.e*.,

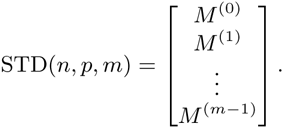

*The columns* 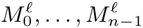 *of each layer are given by cyclic shifts of a base vector*

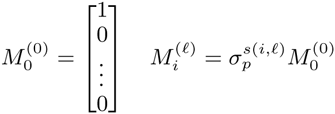

*where* 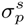 *is a cyclic shift of order s for a vector of length p and*

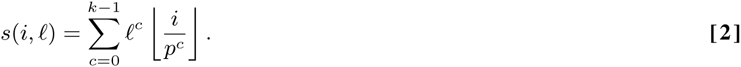

The effect of the cyclic shift 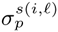 on the vector 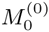 is to shift the 1 from the first position to the *s*(*i*, *ℓ*) mod *p* position in the vector 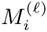.

### Proposition 1

*When q is a prime number p*, *the pooling design M from the* [*m*, *k*, *m*−*k*+1]*_q_-RS code is the same as the STD*(*n* = *p^k^*, *p*, *m*) *(up to the ordering of the columns and the layers)*.

#### Proof

We will start from the definition of the STD matrix and show that the columns of the concatenated layers *M*^(^*^ℓ^*^)^ are exactly the columns of *M*, as defined for Reed-Solomon codes.

We use the convention that a vector of length *p* has components labeled 0 through *p* − 1, then the position of the 1 in the vector 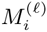 is simply *s*(*i*,*ℓ*) mod *p*. All of the computations in Equation [**2**] can, therefore, be executed modulo *p*. That is, we can replace all of the terms of the form 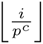 with the expression 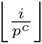 mod *p* and observe that this expression is nothing other than the *c*th digit *b_c_* in the base *p* expansion of *i*,

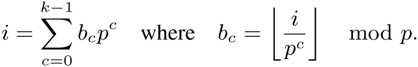

Let *b* =(*b*_0_*,b*_1_,…,*b_k_*_−1_) denote the vector of coefficients in the *p*-ary expansion of *i* and set 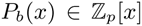 to be the polynomial with coefficients given by the components of *b*

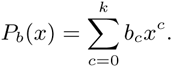

With these definitions, it is clear that *s*(*i*, *ℓ*)= *P_b_*(*ℓ*) and that the position of the 1 in the vector 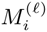 is nothing other than the value of the polynomial *P_b_*(*x*) evaluated at *x* = *ℓ*. Furthermore, there are *p^k^* possible vectors *b* (and hence, *n* = *p^k^*, as desired).

**Combinatorial group testing**. In the traditional combinatorial group testing framework, we express a pooling design by a binary *t* × *n* matrix *M* = (*M_ij_*), where *M_ij_* = 1 if and only if item *j* is present in pool *i*. In the context of this paper, we can think of an item as a specimen; but as we are discussing the fundamentals of group testing in this section, we will continue to use the traditional terminologies for group testing. The *weight* of a row or a column of this matrix is the number of 1’s present in it. The column weight of the *j*’th column of *M* thus represents the number of pools that item *i* belongs, and the row weight of the *i*’th row is the number of items pooled together in pool *i*.

Each pool tests *positive* if there is at least one positive item in the pool. The semantic of positivity depends on the application at hand. The test outcomes of all pools can thus be represented by a vector y = (*y*_1_*,y*_2_,…,*y_t_*) ∈ {0, 1}*^t^*, where *y_i_* = 1 if and only if the *i*’th pool’s test outcome is a positive outcome. The key requirement for a pooling design *M* to be sound is that from the outcome vector y we are able to identify the set of at most *d* positive items, where *d* is an apriori bound on the maximum number of positive items. This worst-case assumption on the maximum number of positive items is in the same spirit as the Hamming formulation in coding theory.

If the pooling matrix *M* allows for unique and correct identification of the positive items, then it is said to be a *d-separable matrix*. It is not hard to show that the matrix *M* is *d*-separable if and only if the Boolean unions of any combination of up to *d* columns of *M* are all distinct, and thus they give rise to distinct test outcome vectors y. The notion of *d*-separability does not indicate *how* we identify the positive items given the test outcome vector y; although in principle we can pre-compute all possible unions of up to *d* columns of *M* and store the results in a huge lookup table with 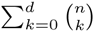 entries.

For moderately large values of *n* and *d*, the lookup table is too large to be useful in practice. Hence, group testing theory has mostly focused on a slightly relaxed notion of pooling design matrix *M* which allows for fast positive item identification. The algorithm for identifying the positive items is very simple: we simply eliminate all items that participate in negative tests, because in case of no testing errors those items are guaranteed to be negative items. This algorithm is called the *naive decoding algorithm*. If the matrix *M* satisfies the property that the naive decoding algorithm always works correctly, i.e. the remaining items are always precisely the positive items, then the matrix is said to be a *d-disjunct matrix*. While it may seem that *d*-disjunctness is a much stronger property than *d*-separability, it turns out that the optimal number of rows of a *d*-disjunct matrix is not too far from the optimal number of rows of a *d*-separable matrix. Hence, by slightly sacrificing the number of tests, we gain a great speedup in decoding time and saving in space requirement for the testing and decoding procedures.

It is also not hard to show that the above algorithmic definition of a *d*-disjunct matrix is equivalent to the following combinatorial definition. The advantage of the combinatorial characterization of disjunct matrices is that it gives us an obvious method to verify whether a given matrix is disjunct.

### Definition 6

(Disjunct Matrix). *A t* × *n binary matrix M is d-disjunct (for* 1 ≤ *d* ≤ *n* − 1*) if and only if the following is true: for any subset* Λ ⊂ [*n*] *of columns such that* |Λ| = *d and an arbitrary column j* ∈ [*n*] − Λ, *there exists a row i such that the jth column has a* 1 *in row i and all columns in* Λ *have a zero in row i*.

As we have alluded to in the introduction, practical applications of group testing often require the pooling matrix to be well-balanced. In particular, it is desirable to have a pooling matrix with close to uniform row weight and close to uniform column weight. The following concept formalizes the notion of a well-balanced matrix.

### Definition 7

(Regular and strongly regular matrix). *A t* × *n binary matrix M is called* (*r*, *c*)-regular *for integers* 1 ≤ *c* ≤ *t and* 1 ≤ *r* ≤ *n if every row has either r or r* + 1 *ones and every columns has either c or c* + 1 *ones. If the rows of the matrix have exactly r ones and the columns exactly c ones*, *we say the matrix is* strongly (*r*, *c*)-regular.

We will show below how to construct disjunct matrices which are (strongly) regular from linear codes. The idea of constructing a disjunct matrix from a linear code dates back to the classic work of Kautz and Singleton on superimposed codes [14]. Superimposed codes are equivalent to disjunt matrices. In fact, Kautz and Singleton already showed us how to construct a disjunct matrix from Reed-Solomon codes, of which the STD design is a special case. The main mathematical contributions of our work is to derive more properties of the RS-code-based construction of disjunct matrices pertaining to practical group testing requirements and to extend this analysis to general linear codes.

### Proposition 2

*Let C be an* [*m*, *k*, Δ]*_q_-linear-code and* 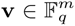 *be an arbitrary vector. Then M*^v+^*^C^ is a strongly* (*q^k^*^−1^*, m*)*-regular matrix that is d-disjunct for any d satisfying the following inequality:*

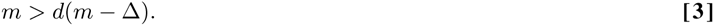

#### Proof

We begin with the case when v = 0. To see that the matrix *M*^v+^*^C^* = *M^C^* is *d*-disjunct, we reason as follows. Let Λ be an arbitrary subset of *d* codewords of *C*, and c an arbitrary codeword not in Λ. Note that there is a one to one correspondence between codewords and the columns of the matrix *M^C^*; hence, we will use “codewords” and “columns of *M^C^*” interchangeably. For each codeword *c*′ ∈ Λ, there are at most *m* − Δ positions *i* ∈ [*m*] for which 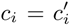. Hence, when *m* > *d*(*m* − Δ) there exists at least one position 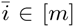 for which 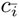 is different from 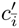 for any codeword *c*′ ∈ Λ. Hence, the row of *M^C^* indexed by the pair 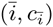 has a 1 in column c and 0 in all columns *c*′ ∈ Λ. We conclude that *M^C^* is *d*-disjunct.

By construction, every column of *M^C^* has exactly *m* ones and by the well known fact that for every *i* ∈ [*m*] and 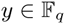, there are exactly *q^k^*^−1^ messages 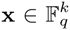 such that *C*(x)*_i_* = *y*, we conclude that every row of *M^C^* has exactly *q^k^*^−1^ ones. This proves that *M^C^* is a strongly (*q^k^*^−1^, *m*)-regular matrix.

Finally, we consider the case when v ≠ 0. In this case *C* still has minimum distance Δ, which allows us to show that *M*^v+^*^C^* is *d*-disjunct exactly as in the argument above for *M^C^*. The column regularity follows by construction and the row regularity follows from the fact that in order to have 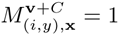 we need *C*(x)*_i_* = *y* − *v_i_*. Thus, we still have the property that for fixed *i* and *y* there are exactly *q^k^*^−1^ messages x such that v + *C*(x)*_i_* = *y*, which implies the desired row regularity.

The last part of the proof shows that when *C* is a *coset* of an [*m*, *k*, Δ]*_q_* code (that is, if 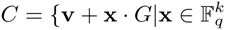 for some 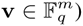), we still get a disjunct design. This property will be useful for generalizing our modularity results. In the main portion of our paper, we observed that RS codes enjoyed modular regularity properties. In fact, this is true of many linear codes as well.

A pooling design matrix *M^C^* constructed directly from an [*m*, *k*, Δ]*_q_*-linear-code *C* has *n* = *q^k^* columns. In practice, we cannot expect the number of items (or specimens) to be an exact power of a prime. Since pooling design matrices constructed from linear codes have very nice properties such as regularity and modularity, we want to retain the construction. The straightforward way out is to select parameters *q* and *k* such that *n* is *just* smaller than *q^k^*, and then remove the last *q^k^* − *n* columns of the matrix. Unfortunately, removing arbitrarily *q^k^* − *n* columns from the matrix *M^C^* might significantly reduce the row-weight uniformity of the matrix, making it unbalanced. To deal with this problem, we impose a stronger property on the linear code *C*. An [*m*, *k*, Δ]*_q_*-linear-code *C* is said to be a *heavy linear code* if there exists a generator matrix *G* for *C* such that the last row of *G* is a vector all of whose components are non-zero. The generator matrix for RS-code shown in Equation 1 has such property. Hence, RS-codes are heavy linear codes. The next proposition explains why heavy linear codes are useful in our context.

### Proposition 3

*Let C be an* [*m*, *k*, Δ]*_q_ heavy linear code. Let the columns of M^C^ be arranged lexicographically by the messages in* 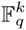*. Then for any positive integer n* ≤ *q^k^*, *consider the matrix M^C^*^|^*^n^ obtained from M^C^ by dropping the last q^k^* − *n columns of M^C^. Then M^C^*^|^*^n^ is an* (⎿*n/q*⏌, *m*)*-regular matrix*.

#### Proof

Since *C* is a heavy linear code, it has a generator matrix *G* such that its last row, which we will denote by g =(*g*_1_,…,*g_m_*), has all non-zero values. Let *G*′ denote the matrix obtained from *G* by removing the last row *vg*. We will use the generator matrix *G* while defining the entries of *M^C^*. We now prove the regularity of *M^C^*^|^*^n^*. The column regularity of *M^C^*^|^*^n^* remains the same as that of *M^C^*. So we only need to be concerned about the row regularity of *M^C^*^|^*^n^*.

Consider the last *q* columns of *M^C^*. Note that they are indexed by the *q* messages of the form (x′*,α*) where x′ is some vector in 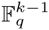 and 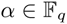 varies over all values in 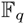. For any 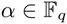, we have

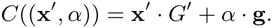

For a fixed position *i* ∈ [*m*], because *g_i_* =0 when *α* varies over all *q* values in 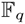 the product *α* · *g_i_* also takes all values in 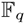. This means the coordinate *C*((x′, *α*))*_i_* varies over all of 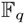 when *α* does. This fact holds true for all positions *i* ∈ [*m*].

Consequently, when we remove the last *q* columns from *M^C^*, we reduce the row weight of *M^C^* by exactly 1. Since the above reasoning holds for any x′, we can continue the argument above untill we are left with *n* columns. If *q* does not divide *n*, then we will remove less than *q* columns in the last chunk of *q* columns. Again by the argument above, it can be seen that the final row weights will either be ⎿*n/q*⏌ or ⎿*n/q*⏌ + 1.

**Modularly disjunct matrices from linear codes**. We have given examples of the modularity of RS designs. In this section, we support and generalize these examples. We formalize the notion of mudularity that is a desired property of pooling matrices as alluded to in the introduction. Informally, we call a matrix *M strictly modularly disjunct* if (a) it itself is a regular disjunct matrix and (b) more importantly, certain sub-matrices are also regular disjunct matrices (but with strictly better disjunctness). We have given examples of the modularity of RS-based designs in the main portion of the paper.

### Definition 8

(Modular matrix). *We say that a t* × *n binary matrix M is* (*d*, *q*)*-modularly disjunct*, *where* 1 ≤ *d* ≤ *n* − 1 *and q divides n*, *if the following hold:*

i. *M is an* (*r*, *c*)*-regular d-disjunct matrix*.
ii. *For* 1 ≤ *i* ≤ *q*, *let* 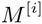 *be the t* × *n/q column submatrix of M that contains the ith contiguous chunk of n/q columns. Then each matrix M*^[^*^i^*^]^ *is an* (*r/q*, *c*)*-regular d*′*-disjunct matrix for some d*′ ≥ *d*.

In fact, we would like to construct such matrices where the modularity property holds for multiple “recursion” levels. We will show in the next section how to construct such nice matrices from linear codes. Towards this end, we define the recursive versions of modularly disjunct matrices.

### Definition 9

(Strictly modular matrix). *Let* 1 ≤ *d* ≤ *d*2 ≤ · ·· ≤ *d_ℓ_* ≤ *n be a sequence of integers such that at least one of the inequalities is strict. Then we call a t* × *n binary matrix M to be* ((*d*_1_,…,*d_ℓ_*), *q*, *ℓ*)*-strictly modularly disjunct (where q divides n)*, *if the following hold. M is a* (*d*_1_*,q*)*-modularly disjunct matrix. Further*, *if ℓ* > 1, *then for every* 1 ≤ *i* ≤ *q*, *M*^[^*^i^*^]^ *is* ((*d*_2_,…,*d_ℓ_*), *q*, *ℓ* − 1)*-strictly modularly disjunct*.

Our main contribution here is to identify two natural properties of linear codes *C* that are sufficient to ensure that *M^C^* is a strictly modularly disjunct matrix. The first property is that in addition to *C* having good distance, certain subsets of *C* (which themselves are linear codes) also have good distances. This is formally captured by the notion of *nested distance* (in the supplementary material). The intuition is that we can apply Proposition 2 to these “sub-codes” to get good disjunctness. The second property is that the linear code has to be *heavy*. This generalizes the property of RS codes we saw previously, where two carefully chosen subsets of columns were equivalent. We formally argue in the supplementary material that a linear code with good nested distance that is heavy results in strictly modularly disjunct pooling designs.

As in the previous section, the order in which we label the columns of *M^C^* makes a difference in the practical application of the design. There are applications where we would like to pool samples in blocks, perhaps at different times, and then store those blocks to be tested jointly or mixed and matched with other populations in different experiments. That is, we would like the columns of the pooling matrix to be modular. We now show how a code with good “nested” distance leads to a strictly modularly disjunct matrix. We first fix a notation, given a *k* × *m* generator matrix *G* and an *i* ∈ [*k*], let *G_i_* denote the matrix obtained from *G* by removing the first *i* − 1 rows of *G*.

### Definition 10

*Let* 1 ≤ Δ_1_ ≤ Δ_2_ ≤ · ·· ≤ Δ*_k_ be integers. Let G be a k* × *m generator matrix for an* [*m*, *k*, Δ_1_] *code C. We say C has a* nested distance *of* (Δ_1_,…, Δ*_k_*) *if for every i* ∈ [*k*], *the code corresponding to G_i_ is an* [*m*, *k* − *i* +1, Δ*_i_*]*_q_-code*.

### Proposition 4

*Let m* ≥ Δ*_k_* ≥ Δ*_k_*_−1_ ≥ ⋯ ≥ Δ_1_ *be integers. For every i* ∈ [*k*], *define d_i_ to be the largest integer such that*

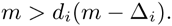

*Then if C is an* [*m*,*k*]*_q_ linear code with nested distance* (Δ_1_,…, Δ*_k_*)*, then M^C^ is a* ((*d*_1_,…, *d_k_)*, *q*, *k*)-*strictly modularly disjunct matrix*.

#### Proof

Since *C* is an [*m*, *k*, Δ_1_]*_q_* linear code, then Proposition 2 implies that *M* is *d*_1_-disjunct and is (*q^k^*^−1^, *m*)-regular, as desired. Let us now consider the *q* submatrices (*M^C^*)^[^*^i^*^]^ for *i* ∈ [*k*]. These matrices correspond to the cosets of the linear code corresponding to the generator matrix *G*_2_, which by definition have distance Δ_2_. This implies that each of these matrices are *d*_2_-disjunct. The proof can then be completed by induction. Next, we fill in the details.

Recall that the columns of *M^C^* are indexed in lexicographic order by the messages in 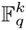. For the rest of the argument fix an *i* ∈ [*k*]. Then note that the columns of (*M^C^*)^[^*^i^*^]^ are indexed by the messages 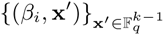, where *β_i_* is the *i*th element in the lexicographic ordering of the elements in 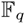. In other words, the columns in (*M^C^*)^[^*^i^*^]^ for 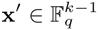 corresponds to the codewords

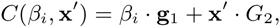

where g_1_ is the first row in *G*. In other words, the codewords corresponding to columns in (*M^C^*)^[^*^i^*^]^ correspond to a coset of the code generated by *G*_2_. Thus, by Proposition 2, (*M^C^*)^[^*^i^*^]^ is *d*_2_-disjunct matrix that is (*q^k^*^−2^, *m*)-regular. Applying this argument inductively (and noting that for any linear code *C* and a coset *C*′, v + *C*′ is also a coset) completes the proof.

Propositions 3 and 4 imply the following result.

### Theorem

*Let m* ≥ Δ*_r_* ≥ Δ*_r_*_−1_ ≥ ⋯ ≥ Δ_1_ *be integers. For every i* ∈ [*r*], *define d_i_ to be the largest integer such that m* > *d_i_*(*m* − Δ*_i_*)*. Suppose C is an* [*m*, *s*]*_q_ heavy linear code with nested distance* (Δ_1_,…, Δ*_k_*), *then for any* 1 ≤ *ℓ* ≤ *r and n which is a multiple of q^ℓ^*, *one can construct an mq* × *n matrix M^C^ that is a* ((*d*_1_,…,*d_ℓ_*), *q*, *ℓ*)*-strictly modularly disjunct matrix*.

**Random linear codes**. We use standard notations employed in computer science to describe asymptotic properties of our codes and pooling designs. Specifically, we denote by *O*(*f*(*n*)) any function that grows at a rate not *faster* than *f*(*n*), as *n* grows (*n* → ∞). Similarly, we denote Ω(*f*(*n*)) a function that grows at a rate not *slower* than *f*(*n*). Finally, we denote by Θ(*f*(*n*)) a function which is both *O*(*f* (*n*)) and Ω(*f* (*n*)), that is, a function growing at the same rate (up-to a constant) as *f*(*n*). More formal definitions can be found in [5].

While our previous result, Theorem 1, explains how to obtain a modularly disjunct matrix from a heavy linear code with the right nested distance properties, it does not explain how to obtain efficiently a code with such properties. In the rest of this section, we show that we can construct a modularly disjunct matrix efficiently from cosets of a random linear code.

Consider a random *k* × *m* matrix *G* over 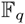. The corresponding linear code *C* is known to satisfy the Gilbert-Varshamov (GV) bound [24, 9, 18]; i.e., its distance is at least 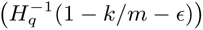 with probability at least 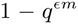, where *H_q_*(*x*) denotes the q-ary entropy function, namely

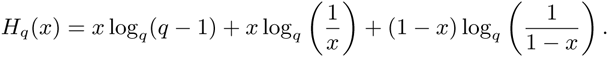

Porat and Rothschild [20] showed that for *q* = Θ(*d*) and *k* = Θ(*m/*(*d* log *d*)), the distance is at least 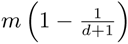. More importantly, they showed how to use the method of conditional expectation [1] to compute the generator matrix of such a code in time *q^O^*^(^*^k^*^)^. Thus, Proposition 2 implies that the corresponding pooling design matrix is *d*-disjunct.

We show that, in addition to being *d*-disjunct and efficiently constructable, a random linear code is also heavy and has good nested distance. To show that a random linear code is heavy, we need the following result.

### Lemma 1

([3]). *Let C be an* [*m*, *k*, Δ]*_q_ linear code and let its dual code have distance* Δ^⊥^ + 1*. Then the number of codewords in C with all non-zero values is lower bounded by*

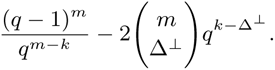

We will use the following two calculations.

### Lemma 2

*Let* 1 ≤ *ℓ* ≤ *k* ≤ *m be positive integers. Let q be an integer such that q* ≥ 2*e*^2^*m/ℓ (where e* =2.71828 *… is the Euler number)*, *then*

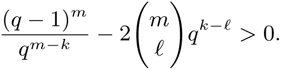

#### Proof

Noting that *q* ≥ 2*e*^2^*m/ℓ* > 8*m/ℓ* > *m/ℓ* +1, and that 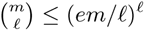, we have

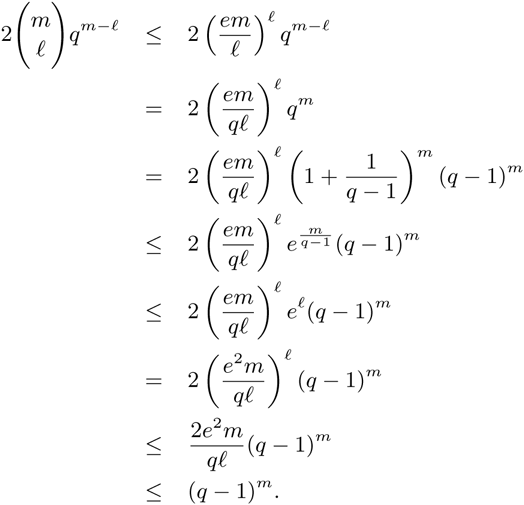

Rearranging the factors gives the desired inequality.

### Lemma 3

*For every integer d*, *let q be an integer such that q* ≥ *cd for a sufficiently large constant c. Then for any x* ≤ 1/2

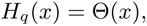

*and for any x* =1 − Θ(1*/d*),

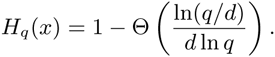

#### Proof

We use the first-order Taylor approximation for sufficiently small real numbers *x* > 0, ln(1 + *x*)= *x* + Θ(*x* ^2^) and ln(1 − *x*)= −*x* + Θ(*x*^2^). Note that since 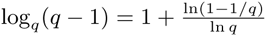, the first term of *H_q_*(*x*) is

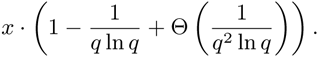

For the last two terms define *y* = min(*x*, 1 − *x*) ≤ 1/2. Since the sum of the last two terms is the same for *x* and 1 − *x*, we just bound their sum in terms of *y*. In this case the second term is

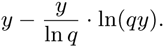

The third term is 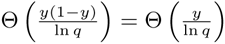.

First, consider the case when *x* ≤ 1/2. In this case *y* = *x* and it can be checked all the terms are dominated by *y*, which implies that *H_q_*(*x*) = Θ(*x*) as required.

Finally, consider the case *x* =1 − Θ(1*/d*). In this case *y* =1 − *x*. In this case, the sum of the first two terms is dominated by the term 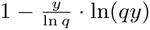. Assuming *q* is at least *c* · *d* for large enough *c*, the 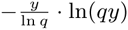 term dominates the third term, which implies the claimed bound.

Consider a [*m*, *k*, (1 − 1/(*d* + 1))*m*]*_q_* random linear code. By Proposition 2, the corresponding matrix will be *d*-disjunct. Let us now figure out the parameters *q* and *k* in terms of *m* and *d*. It is well-known that the random code lies on the GV bound which by Lemma 3 implies that

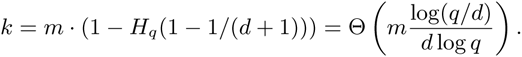

It is also well-known that with high probability the dual of this code lies on the GV bound; i.e., the dual distance satisfies

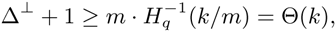

where the equality follows from Lemma 3. Further, it is easy to show that the dual distance can be at most *k* (this follows e.g. from the Singleton bound). Thus, we have

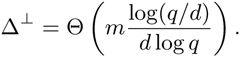

We now want to use Lemmas 2 and 1 to imply that the random code with high probability is heavy. To do this we need

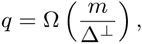

or

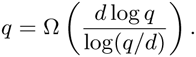

It is not too hard to check that the above is true if we pick

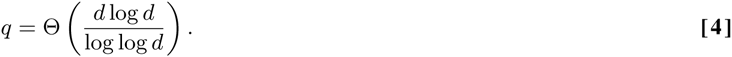

This along with the restrictions above implies that we have

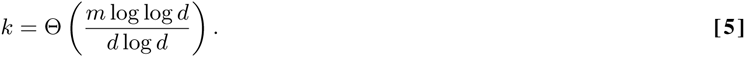

The corresponding matrix will have *m* · *q* rows, which, by recalling that *n* = *q^k^*, leads to 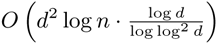 rows.

**Good Nested Distance**. We now observe that the cosets of a random linear codes have good distance. In particular, note that for a random *G*, the matrix *G_i_* is also a random matrix (though with smaller number of rows). Thus, by the known results with high probability, all these codes lie on the GV bound.

There are a couple of caveats in trying to combine the two results on heaviness and nested distance: First, while calculating the distance of the cosets, we cannot go down to *G_k_* because the high probability bound is not small enough to take a union bound over all the values of *i* ∈ [*k*]. Second, even though a random linear code is heavy with high probability, it is not necessary that one of the rows of the random *G* will have all non-zero values. However, we can always recompute another generator matrix *G*′ such that its last row is all non-zeroes. In fact, one can argue that one can obtain *G* from *G*′ by putting the all non-zero row at the bottom of *G* and then removing one of the *k* rows of *G*.^1^ Again, while making the argument for the dual distance to be too large we cannot go down to small sub-matrices *G_i_*. Further, to make the argument for showing that a random linear code is heavy, we need Δ^⊥^ = Θ(*m*/*d*), which implies that we can only consider *G_i_* for *i* ≤ *ℓ* = *O*(*k*).

The above two points imply that we can only only “recurse” up to say *k*/2 levels (instead of going all the *k* levels to go down to *G_k_*). This means we will obtain a strictly modularly disjunct matrix with Θ(*k*) levels of recursion, which for practical applications such as ours is fine.

Next, we quickly outline how one can adapt the algorithm of [20] to our case above. Note that what we need is the following for the linear codes corresponding to the generator matrices *G_k_,G_k_*_−1_,…,*G_O_*_(_*_k_*_)_: both the code and its dual has to lie on the GV bound. In particular, we have to run *O*(*k*) “copies” of the argument in [20]. However, this will not affect the final *q^O^*^(^*^k^*^)^ run time, which implies that

### Theorem 2

*One can construct a* ((*d*_1_,…,*d_ℓ_*), *q*, *ℓ*)*-strictly modularly disjunct matrix (with ℓ* = Θ(*k*)*) with the* 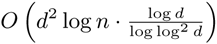 *number of rows in time polynomial in n*, *where q and k are chosen as in (4) and (5)*.

**Computing the average disjunctness of Reed-Solomon Codes**. In this section, we address the average disjunctness of Reed-Solomon codes, and in particular the problem of computing it. As mentioned in the main body of the paper, disjunctness, which is a worst-case guarantee, is often too strong in practice. To that end, we consider *average disjunctness*.

### Definition 11

((*δ*, *d*)-Disjunct Matrix). *A t* × *n binary matrix M is* (*δ*, *d*)*-average-disjunct* (*for* 1 ≤ *d* ≤ *n* − 1) *if and only if the following is true. Let* Λ ⊂ [*n*] *be a set of d columns chosen uniformly at random. Then with probability at least* 1 − *δ*, *for any column j* ∈ [*n*] − Λ, *there exists a row i such that the jth column has a* 1 *in row i and all columns in* Λ *have a zero in row i*.

We first observe that the average disjunctess can be bounded in terms of the roots of sets of polynomials.

### Proposition 5

*Let* Ω *be a set of polynomials of degree at most k over* 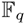, *so that* |Ω| = *n*, *and further so that* Ω *forms an additive subgroup (i.e. every p*_1_*,p*_2_ ∈ Ω, *we have p*_1_ ± *p*_2_ ∈ Ω*). Let* 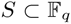*. For a subset* Λ ⊂ Ω, 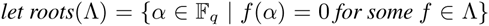. *Let N_d_*(*S*, Ω) *be the number of sets* Λ ⊂ Ω \ {0} *of size d so that S* ⊂ *roots*(Λ)*. Then for any δ so that*

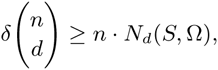

*RS*(*S*, Ω) *is* (*δ*, *d*)*-average-disjunct*.

#### Proof

Say that Λ is *bad for p* ∈ Ω (with respect to *S*) if for all *α* ∈ *S*, there is some *f* ∈ Λ so that *f*(*α*) = *p*(*α*), and similarly that Λ is *bad* if there exists a *p* for which it is bad. For a fixed *S*, Ω, let 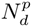 be the number of size *d* sets Λ ⊂ Ω, not containing *p*, so that Λ is bad for *p* with respect to *S*. Thus, the number of sets Λ that are bad is at most 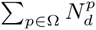. By definition, *RS*(*S*, Ω) is (*δ*, *d*)-average-disjunct for any *δ* so that

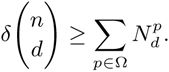

We will show that in fact 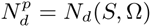 for all *p*, and this will complete the proof. First, notice that by definition, 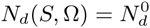. Further, if Λ is bad for 0, then Λ+ *p* = {*f* + *p* | *f* ∈ Λ} is bad for *p*, and conversely if Λ is bad for *p*, then Λ − *p* is bad for 0. Thus, the sets Λ which are bad for 0 are in bijection with those bad for *p*, for all *p*, which completes the proof.

Next, we address the issue of computing the average disjunctess of Reed-Solomon codes. In order to compute the results reported in Figure 5, we use Proposition 5. Recall that in Figure 5, Ω is the set of Reed-Solomon codewords corresponding to constant, linear, and monic quadratic polynomials, and that *S* is an arbitrary subset of 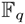 of size *m* = 6. We must compute the number of subsets Λ ⊂ Ω of size *d*, so that the roots of the polynomials in Λ cover *S*. Our argument below will show that in fact this number is independent of the choice of *S*, and so our bounds hold for all choices of layers.

Rather than computing the number of bad sets Λ, we will equivalently compute the probability that a random set Λ of size *d* is bad. We may do this recursively. Let *p*(*m*, *r*, *t*) denote the probability that *r* polynomials cover an arbitrary set *S_t_* of size *t*, where the polynomials are drawn uniformly at random from an any set Ω*_m_* of size *m* so that Ω*_m_* ⊂ Ω and

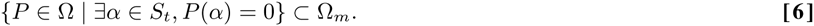

We note that by symmetry, *p*(*m*, *r*, *t*) is well defined—that is, that it does not depend on the choice of *S_t_* or Ω*_m_*, as long as they are compatible in the sense of [**6**].

Then *p*(*m*, *r*, *t*) obeys the recursive relationship

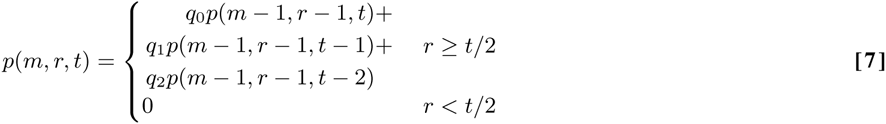

where *q_i_* is the probability that a randomly chosen polynomial from Ω*_m_* has exactly *i* roots in *S_t_*. Indeed, if *r* < *t*/2, then there is no way that *r* polynomials of degree at most two can have between them *t* roots. On the other hand, if *r* ≥ *t*/2, then there are three cases: either the first polynomial includes no roots of *S_t_*, one root of *S_t_*, or two roots of *S_t_*, and these are the three terms in the sum.

Because the number of polynomials in Ω*_m_* with precisely two roots in *S* is 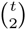 (indeed, this is the number of such polynomials in Ω, and all of them are contained in Ω*_m_* because of [**6**]), we have

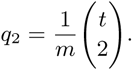

Similarly,

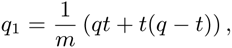

because the polynomials in Ω that have precisely one root in *S_t_* are of the form *β*(*x* − *α*) or (*x* − *α*)^2^ for *α* ∈ *S_t_*, 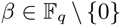, or of the form (*x* − *β*)(*x* − *α*) for *α* ∈ *S_t_*, *β* ∈ *F_q_* \ *S_t_*. Finally,

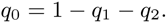

Using [**7**], we may easily compute

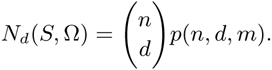

**Blue Dye Decoding**. We combined 480 samples (150*µ*l of water or 100X diluted blue dye) into 96 pools, using the Reed-Solomon design with the following parameters:

- source volume = 120*µ*l
- each step = 20*µ*l
- basic window = 16
- weight =6
- specimens per pool = 30
- offset =0

The actual well volume should be at least

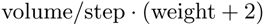

to account for pipetting inaccuracy and to prevent specimen dropout. The pooling was performed using a Tecan Evo liquid-handling robot. After pooling (≈ 15 hours), 100 *µ*l aliquots were manually transferred from the deepwell pooling plate to a clear 96 well flat bottom plate in order to measure absorbance. (The reference wavelength (emission) was 470nm; and the measurement wavelength (excitation) 630nm.)

We decoded the results using GPSR [19] using the absorbance measurements for the 96 pools. We adjusted *τ* (a tuning parameter) to balance noise and sparsity.

**HapMap Decoding**. We took 96 samples from the International HapMap Project and normalized them to 20ng/*µ*l. Then we pooled them into 25 pools using Shifted Transversal Design with the following parameters:

- source volume = 100*µ*l
- each step = 20*µ*l
- basic window = 5
- weight = 5
- specimens per pool = 96/5
- offset = 0

The 25 pools were barcoded and enriched by hybrid selection using Agilent SureSelect baits from the Framingham Heart Study ( 800kb). The pools were further compressed into five mega pools and sequenced on five lanes on the Illumina HiSeq to 1000X average coverage per pool. Reads were aligned with BWA [16] and bam files were separated by barcode. Variants were called with Varscan [6] with the following parameters: `mpileup2cns --min-var-freq 0.00001 --min-reads2 1 --p-value 0.99`. Duplicate reads were not removed as we found that doing so diminishes the concordance between Sudoku calls and 1000 Genomes.

Indels and variants on the X chromosome were excluded and if found in more than five samples (by sequencing or in 10000 Genomes). Only variants with minimal coverage 3000X and passing all filters were kept for analysis. We compared calls between 75 of our samples that were also sequenced as part of the 1000 Genomes Project, focusing on autosomal SNPs with MAF ≤ 2%.

## Footnotes

1 Start from the bottom row of *G* and greedily remove the first row in *G* that in augmented matrix is no longer independent of the rows below it.

